# Restoring transient connectivity during development improves dysfunctions in fragile X mice

**DOI:** 10.1101/2024.09.08.611918

**Authors:** Dimitri Dumontier, Samuel A. Liebman, Viet-Hang Le, Shanu George, Deasia Valdemar, Linda Van Aelst, Gabrielle Pouchelon

## Abstract

Early-generated circuits are critical for the maturation of cortical network activity and the formation of excitation/inhibition (E/I) balance. This process involves the maturation of specific populations of inhibitory neurons. While parvalbumin (PV)-expressing neurons have been associated with E/I impairments observed in neurodevelopmental disorders, somatostatin-expressing (SST) neurons have recently been shown to regulate PV neuron maturation by controlling neural dynamics in the developing cortex. SST neurons receive transient connections from the sensory thalamus, yet the implications of transient connectivity in neurodevelopmental disorders remain unknown. Here, we show that thalamocortical connectivity to SST neurons is persistent rather than transient in a mouse model of Fragile X syndrome. We were able to restore the transient dynamics using chemogenetics, which led to the recovery of fragile X-associated dysfunctions in circuit maturation and sensory-dependent behavior. Overall, our findings unveil the role of early transient dynamics in controlling downstream maturation of sensory functions.

## Introduction

The balance between excitation and inhibition (E/I) is critical for cortical network activity and sensory encoding. The E/I ratio is formed during development with the progressive integration of specific GABAergic inhibitory interneurons into functional network motifs. In adulthood, parvalbumin-expressing (PV) interneurons provide feedforward inhibition in response to sensory thalamic activity, while somatostatin-expressing (SST) neurons mediate feedback inhibition^1–3^. Recent evidence indicates that SST neurons transiently receive sensory thalamocortical (TC) inputs during early postnatal development^4–6^. This process is involved in early neural dynamics and controls the integration of PV neurons in cortical circuits^7–11^, which is essential for the development of adult mouse behaviors^12^. Although transient connectivity controls critical processes of cortical development, the implications in neurodevelopmental disorders remain unexplored.

E/I network imbalance, resulting in cortical hyperexcitability, has long been implicated in the etiology of neurodevelopmental disorders, such as autism spectrum disorders, including fragile X syndrome (FXS)^13–16^. Hyperexcitability of sensory upper-layer cortical circuits leads to hypersensitivity and tactile processing abnormalities, which are common to most autism disorders, including FXS^17,18^. As the main targets of sensory inputs in adults, PV neurons have been extensively investigated. They are thought to underlie hypersensitivity associated with FXS and other neurodevelopmental disorders^17,19–22^. However, these neurons mature only after critical periods of development, while FXS-associated hypersensitivity is observed at earlier postnatal stages^23–25^. In contrast, early activity of SST neurons, driven by sensory TC inputs, suggest that transient TC connectivity to these neurons orchestrate cortical maturation and play a critical role in the etiology of neurodevelopmental disorders. The developing programs of SST neurons could therefore be an ideal therapeutic target in the context of early intervention. However, transient connectivity to SST neurons in neurodevelopmental disorders remains unexplored to date.

Here, we show that somatosensory TC inputs to SST neurons, which are normally transient, are persistent in a mouse model of FXS, the *Fmr1* knockout (KO). Chemogenetic manipulation of SST neurons within a critical time window regenerates the transient dynamics of the TC circuit. This restoration of transient TC inputs to SST neurons alleviates *Fmr1* KO sensory impairments observed within cortical circuits and tactile behaviors. Thus, SST neurons involved in transient TC connectivity are an essential developmental trigger of subsequent circuit maturation and processing of sensory inputs.

## Results

### TC connectivity to SST neurons is persistent rather than transient in *Fmr1* KO mice

In order to determine whether transient connectivity to SST neurons is involved in FXS, we first performed a longitudinal analysis of SST cell responses to TC inputs in the primary somatosensory cortex (S1) during development (postnatal day (P)5-P40) in control and *Fmr1* KO mice, the most common model of FXS^26^ (Figure 1A). SST neurons were identified using GFP reporter expression following AAV-DIO-GFP injection at P0 (Fig. 1A) in SST^Cre^::WT and SST^Cre^::*Fmr1* KO animals. SST neuron responses in layer 5 (L5) of S1, where they are known to form transient connections, were recorded upon electrical stimulation of TC fibers in TC-slices prepared for each age window (Figure 1B-C; Figure S1A-B. Early P5-P6; Intermediate P10-P16; Late P30-P40)^2,27,28^.

**Figure 1:**
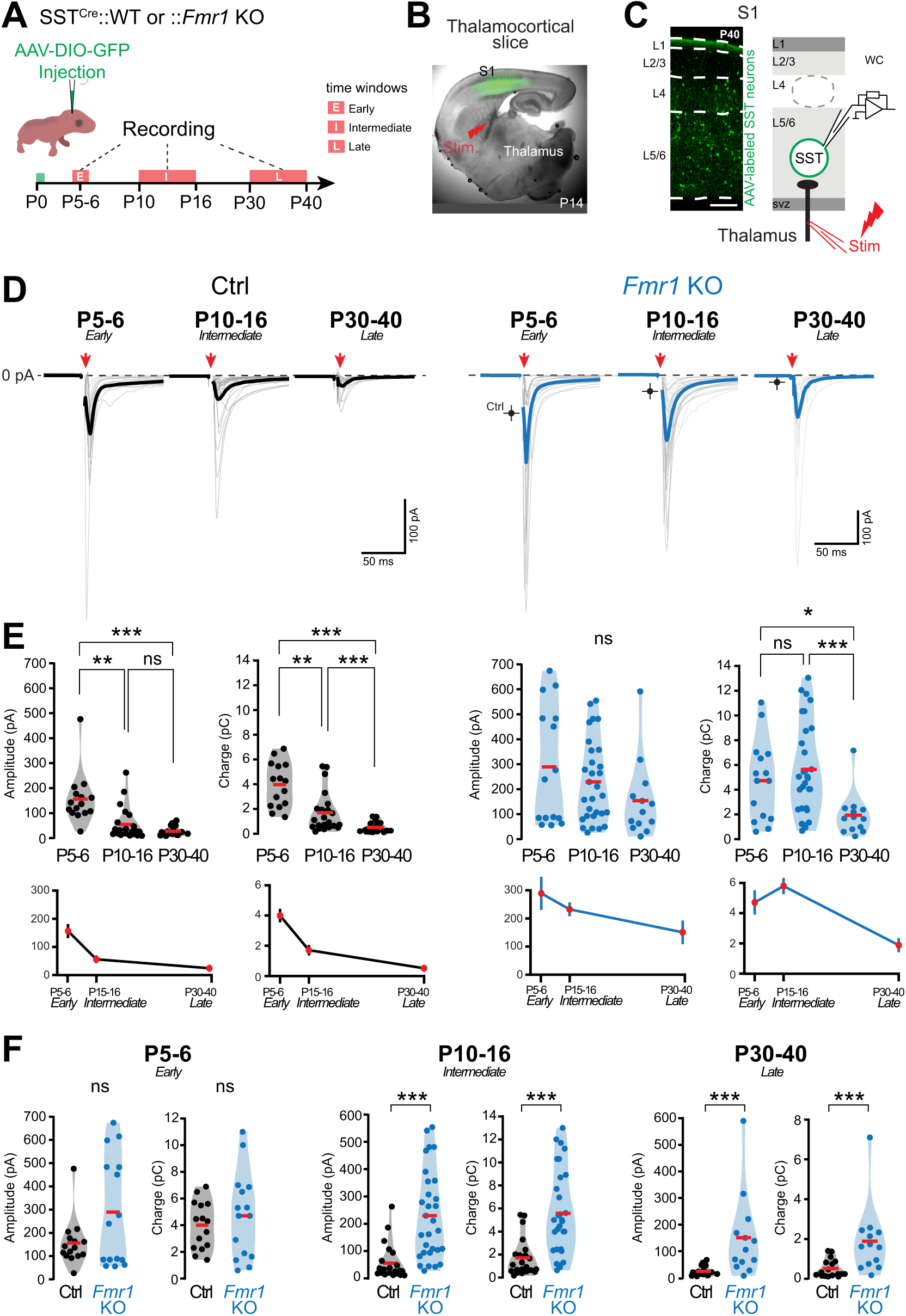
TC connectivity to SST neurons is persistent rather than transient in *Fmr1* KO mice. **A**) Experimental strategy and timeline of AAV-DIO-GFP injection at P0-1 to label SST neurons within control SST^Cre^::WT and SST^Cre^::*Fmr1* KO animals, and recording during three time windows: Early (E) P5-P6; Intermediate (I) P10-P16; Late (L) P30-P40. **B**) Example of TC slice expressing the AAV-DIO-GFP in S1 SST neurons at P14. Stim = electrical stimulation of TC fibers. **C**) Example of SST neurons labeling within cortical plate of S1 and schematic illustration of SST neuron recordings. Scale bar:200 μm. **D**) Averaged EPSCs evoked in every L5 SST cell by TC electrical stimulation (gray) and their population average from control (black) and *Fmr1* KO (blue) animals, showing the transient dynamics versus persistence of TC inputs onto SST neurons. Black and blue dots represent the populational EPSC peak Ctrl and *Fmr1* KO. **E**) Amplitude and charge of the averaged EPSCs evoked in L5 SST cells from P5 to P40 in Ctrl (Ctrl; Amplitude; Early (E: P5-6): 156.14 ± 25.1 pA, n = 15, nSL = 3, N = 2F/1M; Intermediate (I: P10-16): 55.79 ± 13.56 pA, n = 22, nSL = 12, N = 3F/3M; Late (L: P30-40): 24 ± 4.4 pA, n =17, nSL = 4, N = 2F/2M; Kruskal-Wallis test p=4.09e-6; E/I p=4.7e-4, I/L p=0.08, E/L p=9.6e-7; Dunn’s post hoc tests; Charge; E: 3.99 ± 0.44 pC; I: 1.7 ± 0.33 pC; L: 0.51 ± 0.1 pC; p=3.54e-7; E/I p=0.0015, I/L p=0.0068, E/L p=5.06e-8) and *Fmr1* KO (KO; Amplitude; E: 289.2 ± 58.8 pA, n = 15, nSL = 3, N = 2F/1M; I: 229.74 ± 29 pA, n = 30, nSL =15, N = 6F/3M; L: 150.8 ± 42 pA, n = 13, nSL = 4, N = 3F/1M; Kruskal-Wallis test p=0.175; Charge; E: 4.71 ± 0.8 pC; I: 5.58 ± 0.66 pC; L: 1.88 ± 0.46 pC; p=0.0015; E/I p=0.50, I/L p=3.7e-4, E/L p=0.011). Bottom: Representation of time dynamics with average EPSCs ± SEM across the time windows. **F)** Average EPSC amplitude and charge between Ctrl and *Fmr1* KO animals (Amplitude: E p=0.648, I p=2.02e-6, L p=0.00043; Charge: E 0.739, I p=1.99e-5, L p=0.0011; Bilateral Mann Whitney U test). N = animal, nSL = slice, n = cell replicates. Data are presented as mean ± SEM.

As expected in control animals, SST neuron responses were the strongest during the early window, dropped at the intermediate stage (-64%) and further diminished at the late stage (-85%; Figure 1D). In contrast to control dynamics, SST neuron responses did not weaken after the early stage in the *Fmr1* KO animals (Intermediate: - 21%, Late: -48%; Figure 1D-E). While responses between control and *Fmr1* KO animals were similarly strong during the early stage (P5-6), they persisted throughout the intermediate and late stages in *Fmr1* KO mice (P10-16 / P30-40; Figure 1E-F). Of note, responses from both hemizygous (*Fmr1*-/y) males and heterozygous (*Fmr1*+/-) females, which display mosaic expression of FMRP (Figure S1E)^29^, were not significantly different (Figure S1F-H). The persistence in TC connectivity was specific to SST neurons, rather than a global increase of TC inputs, as neighboring pyramidal cells responses to the same TC inputs were indistinguishable between control and *Fmr1* KO mice at P10-16 (Figure S1C-D). Overall, we identified the persistence of normally transient TC connectivity to SST neurons as a marker of FXS.

### Postnatal chemogenetic manipulation of SST neurons restores the transients in *Fmr1* KO

The observation that normally transient TC connectivity onto SST neurons is persistent in *Fmr1* KO animals prompted us to determine whether restoring these transitory dynamics corrects somatosensory impairments in FXS mice. As previously shown, transient TC connectivity to SST neurons is controlled by metabotropic signaling and can be modulated with chemogenetic manipulations, using Designer Receptors Exclusively Activated by Designer Drugs (DREADD)^30^. Strikingly, sustained activation of DREADD-Gq receptors promotes the premature weakening of TC inputs to SST neurons^12^. Therefore, we employed this approach to regenerate transient dynamics in *Fmr1* KO animals, which we termed the ‘*Rescue’* condition. We injected AAV-DIO-hM3q-mCherry in SST^Cre^::WT and SST^Cre^::*Fmr1* KO mice at birth. Daily injections of Clozapine-N-Oxide (CNO) were administered from P6 to P16, spanning the early and intermediate stages, when TC inputs onto SST neurons normally mature, to mimic the physiological transient pattern (Figure 2A; Figure S2A-B).

**Figure 2:**
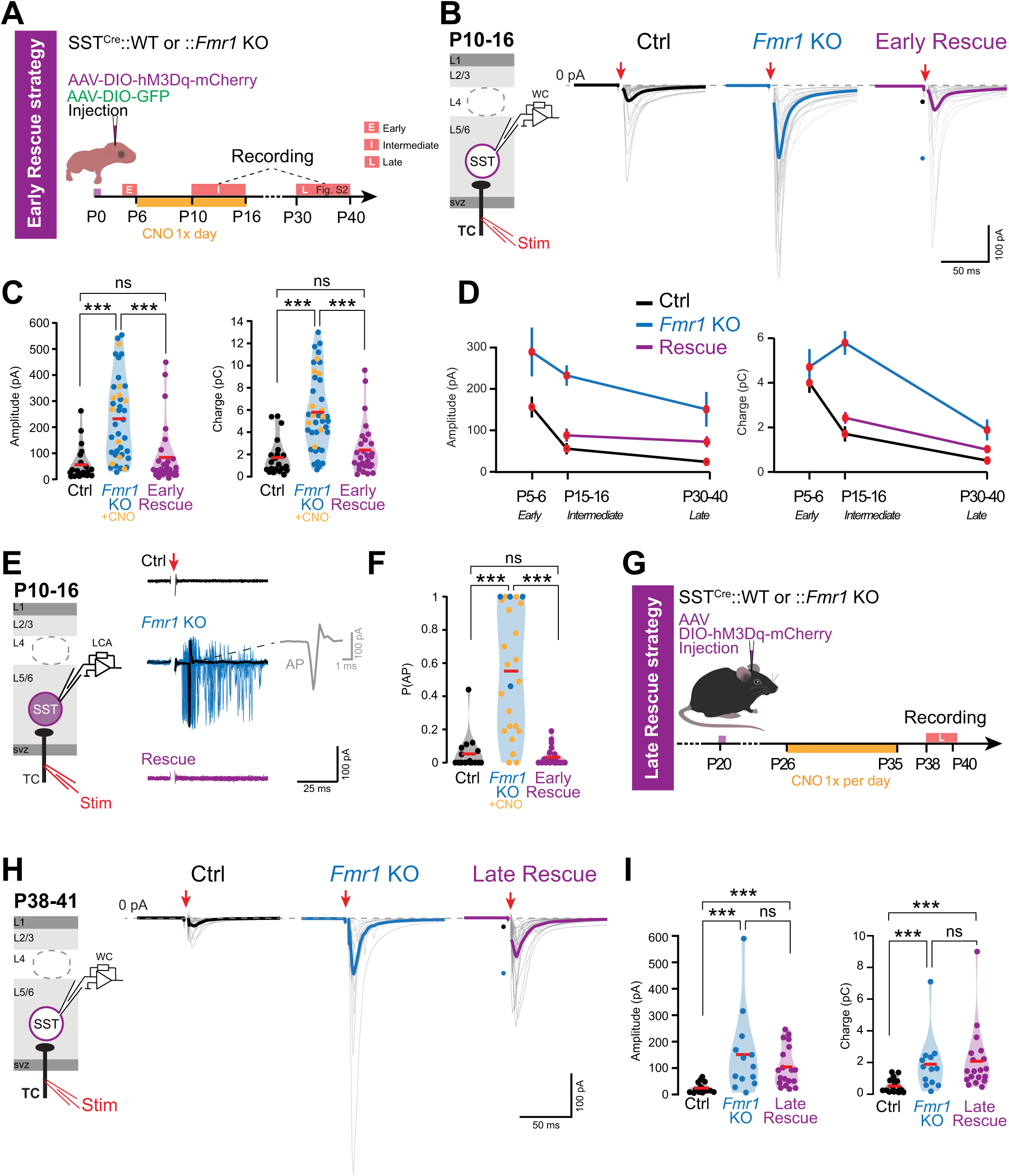
Postnatal chemogenetic manipulation of SST neurons restores the transients in *Fmr1* KO mice. **A**) Schematic and timeline of Early Rescue strategy: P0-1 injection of AAV-DIO-hM3Dq-mCherry in SST^Cre^::*Fmr1* KO mice and AAV-DIO-GFP in SST^Cre^::WT and SST^Cre^::*Fmr1* KO animals for control and early Rescue conditions. Daily injections of CNO right after the Early time window (E) from P6 to P16 maximum. Recordings of SST neurons during intermediate (I) and late (L) time windows. **B**) Average EPSCs evoked in every L5 SST cells by TC electrical stimulation (gray) and their population average from Control (black), *Fmr1* KO (blue) and Early Rescue (purple) animals, during the Intermediate window (P10-P16), showing the restoration of the transient TC input dynamics. Black and blue dots represent the population average EPSC peak of Ctrl and *Fmr1* KO animals. **C**) Amplitude and charge of the average EPSCs during the intermediate time window. Ctrl dataset from Fig.1. *Fmr1* KO dataset comprises data from Figure 1, together with added cells from KO injected with CNO (n = 10, yellow dots. Amplitude: Ctrl: 55.79 ± 13.56 pA, n = 22; KO: 232.48 ± 24.7 pA, n = 40, nSL = 18, N = 8F/4M; Early Rescue: 84 ± 21 pA, n = 29, nSL = 8, N = 2F/6M; Kruskal-Wallis test p = 1.86e-8; Dunn’s post hoc tests: Ctrl/KO p =1.31e-07, KO/Early Rescue p = 4.09e-06, Ctrl/Early Rescue p=0.327. Charge: Ctrl: 1.7 ± 0.33 pC, n = 22; KO: 5.79 ± 0.53 pC, n = 40; Early Rescue: 2.38 ± 0.42 pC, n = 29; Kruskal-Wallis test p = 8.5e-8; Dunn’s post hoc tests: Ctrl/KO p = 3.63e-07, KO/Early Rescue p = 1.36e-05, Ctrl/Early Rescue p=0.305). **D**) Amplitude and charge of EPSCs evoked in SST neurons within Ctrl, *Fmr1* KO and Rescue conditions plotted as a function of time windows. **E**) Loose cell-attached recording of SST neuron firing upon TC fiber stimulation. Illustrative example traces of SST cell firing. Inset: high magnification of an action potential (AP). **F**) Firing probability (Ctrl: 0.052 ± 0.025, n = 17, nSL = 4, N = 1F/3M; KO: 0.55 ± 0.07, n = 23, nSL = 3, N = 1F/2M; Early Rescue: 0.032 ± 0.01, n = 24, nSL = 4, N = 3F/1M; Kruskal-Wallis test p = 7.53e-08; Dunn’s post hoc tests: Ctrl/KO p = 4.13e-06, KO/Early Rescue p =3.39e-07, Ctrl/Early Rescue p = 0.887). **G**) Schematic and timeline of Late Rescue strategy: P20 injection of AAV-DIO-hM3Dq-mCherry in SST^Cre^::*Fmr1* KO mice. Daily injections of CNO after the intermediate (I) time window from P26 to P35. Recordings of SST neurons during the late time window (P38-P41). **H**) Average EPSCs evoked in every L5 SST cell by TC electrical stimulation (gray) and their population average from Control (black), *Fmr1* KO (blue) and Late Rescue (purple) animals, showing no effect of Late Rescue manipulation on TC input dynamics. Black and blue dots represent the population average EPSC peak of Ctrl and *Fmr1* KO animals. **I**) Amplitude and charge of the average EPSCs shown in I). Ctrl and *Fmr1* KO datasets from Fig.1 (Late Rescue Amplitude: 106.44 ± 17 pA, n=19, nSL=6, N=3M/2F; Kruskal-Wallis test p=2.68e-5; Dunn post-test, Ctrl/KO p=9.7-5, Ctrl/Late Rescue p=6.6e-5, KO/Late Rescue p=0.77; Late Rescue Charge: 2.07 ± 0.44 pC; Kruskal-wallis test p=7.8e-5; Dunn’s post hoc tests, Ctrl/KO p=8.0e-4, Ctrl/Late Rescue p=5.4e-5, KO/Late-Rescue p=0.754). N= animal, nSL = slice, n= cell replicates. Data are presented as mean ± SEM.

The manipulation of Gq pathways within SST neurons in S1 of *Fmr1* KO animals significantly reduced the strength of TC connectivity during the intermediate (Figure 2B-C) and the late (Figure S2C-D) stages. This approach led to a substantial restoration of the transient dynamics in *Fmr1* KO animals similar to those of control mice during the intermediate window (Figure 2D) without substantial changes in SST neuron intrinsic properties (Table S1). As with *Fmr1* KO observations, the *Rescue* condition showed no significant differences between *Fmr1-*/y and *Fmr1*+/- animals (Figure S2E-F). Transient connectivity from TC inputs to SST neurons in L5 of S1 is essential for downstream circuit maturation involving L5 and L4 E/I balance^4,5,11^. This process is typically regulated by SST neurons, which transition from being active to unresponsive to direct sensory inputs after the first postnatal week^7,31,32^. We therefore next examined whether the persistence of TC connectivity in *Fmr1* KO mice influences L5 neuron activity upon TC fiber stimulation. During the intermediate time window, while TC inputs failed to recruit SST neurons in control mice, they evoked a robust firing of the SST cells in the *Fmr1* KO animals. This activity was abolished when transient TC connectivity was restored in the *Rescue* condition (Figure 2E-F; Figure S2G). In addition, while excitatory responses of L5 pyramidal neurons to TC input stimulation were similar across all conditions, the variance of the inhibitory responses was significantly increased in *Fmr1* KO mice, but restored in the *Rescue* animals (Figure S2H-J).

As transient connectivity typically occurs only during an early critical time window, we next tested whether chemogenetic manipulation of SST neurons later in development restores the TC connectivity dynamics in *Fmr1* KO animals. AAV-DIO-hM3Dq-mCherry was injected at P20, and CNO was administered during the late time window, from P26 to P35, matching the total duration used for early *Rescue* (Figure 2G; Figure S2K). Remarkably, the late *Rescue* did not weaken TC connectivity (Figure 2H-I; Figure S2K-M). Altogether, our results show that chemogenetic manipulation of SST neurons restores the transient dynamics of TC connectivity in *Fmr1* KO mice during a critical time window, when this connectivity normally matures.

### Early restoration of the transient dynamics corrects downstream cortical maturation and tactile dysfunctions in adult *Fmr1* KO mice

Hypersensitivity described in children with FXS has been linked with upper layer (L2/3-L4) hyperexcitability^33,34^ and defects in PV neuron maturation^19,20^. Notably, SST neurons control PV and L4-excitatory neuron circuit maturation during the early postnatal stages of development. We therefore investigated whether transient connectivity regulates S1 circuit integration and whether defects in early dynamics underlie FXS-associated dysfunctions at mature stages (Figure 3A). We first examined the global input integration onto L4 neurons in S1. We recorded L4 neuron firing activity following electrical stimulation of subventricular zone (SVZ) fibers within the same cortical column (Figure 3B-C). L4 neuron firing resulted from local integration of electrically evoked currents as bath application of GABAergic and glutamatergic transmission blockers promoted or abolished L4 neurons firing, respectively (Figure S3A-D). While L4 neuron firing in *Fmr1* KO mice was less robust compared to cells in control animals, the firing probability of L4 cells in the *Rescue* condition was increased to control levels (Figure 3D).

**Figure 3:**
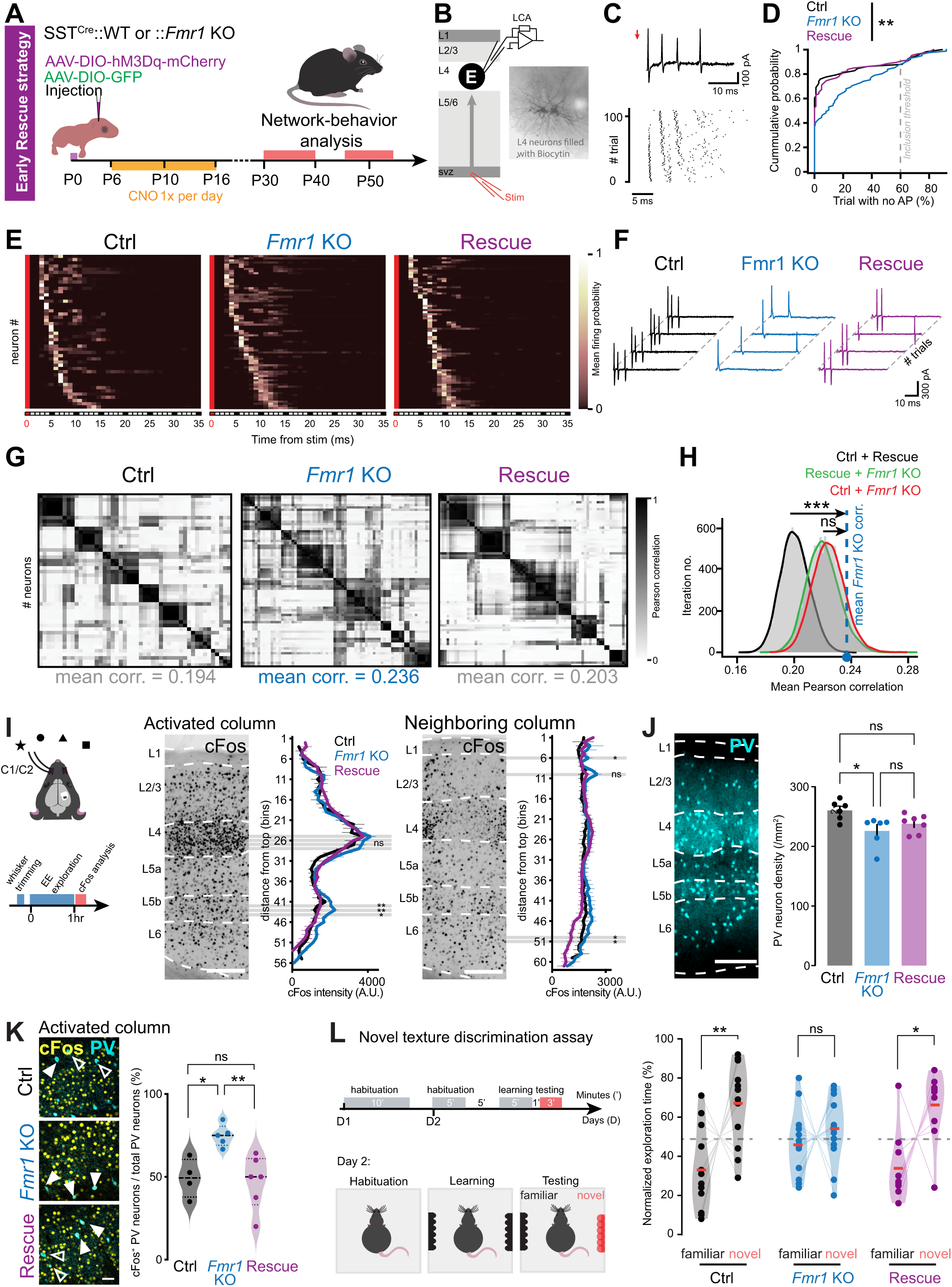
Early restoration of the transient dynamics corrects downstream cortical maturation and tactile dysfunctions in adult *Fmr1* KO mice. A) Schematic and timeline for downstream effect analysis of Early Rescue strategy. **B**) Loose cell-attached recording of L4-excitatory neuron (E) firing upon electrical stimulation of afferents originating from local subventricular zone (SVZ). Inset: morphology of two L4 neurons. **C**) Illustrative example of a control L4 neuron responses to local SVZ electrical stimulation. Top: individual trial. Bottom, raster plot of 110 trials. **D**) *Fmr1* KO L4 neuron failure rate is restored in the Rescue condition. Cumulative probability distribution of the proportion of trials without action potential (AP) (Ctrl: 12.25 ± 3.2, n = 59, nSL = 13, N = 2F/5M; KO: 20.59 ± 2.8, n = 82, nSL = 14, N = 2F/3M; Rescue: 12.24 ± 2.47, n = 87, nSL = 12, N = 5F/4M; Krukal-Wallis test p = 0.007; Dunn’s post hoc tests: Ctrl/KO p = 0.0038, Ctrl/Rescue p = 0.495, KO/Rescue p = 0.014). **E**) Representation of L4-excitatory neurons (E) firing pattern upon SVZ stimulation: Average firing probability per neuron (row) binned at 1 ms in the 35 ms following stimulation (Ctrl: n = 48, nSL = 13, N = 2F/5M; KO: n = 72, nSL = 14, N = 2F/3M; Rescue: n = 74, nSL = 12, N = 5F/4M). Firing patterns are more discrete in the Ctrl and Rescue conditions compared to *Fmr1* KO mice, where they are more dispersed. **F**) Individual neuron firing examples (4 trials) illustrating the dispersed firing pattern of *Fmr1* KO SST neurons. **G**) Correlation matrices of the mean firing probability (from D), showing a discrete organization of Ctrl and Rescue L4 neurons compared to the *Fmr1* KO cells, which self-organize more continuously. **H**) Bootstrap analysis of the matrix correlation data inputs indicates that the values shown in G) are not artifacts of sampling and that only *Fmr1* KO SST neurons trigger an increase in the mean correlation within pooled datasets. The distributions of correlations from the bootstrapped datasets pooled with *Fmr1* KO neurons (‘Rescue+*Fmr1* KO’ and ‘Ctrl+*Fmr1* KO’) are shifted towards the observed mean correlation value of *Fmr1* KO neurons. In contrast, ‘Ctrl+Rescue’ dataset did not recapitulate this shift. The probability of observing a ‘Ctrl+Rescue’ correlation distribution as close to the observed *Fmr1* KO mean correlation is p = 0.00049; ‘Ctrl+ *Fmr1* KO’ p = 0.123; ‘Rescue+*Fmr1* KO’ p = 0.075. **I**) Left: experimental design for cFos expression analysis upon C1/C2 whisker stimulation from enriched environment (EE) exploration. Right: Examples of cFos staining in an activated and a neighboring column of a control animal. Scale bar: 200 μm. The mean intensity across the column shows an increase in cFos expression in *Fmr1* KO animals compared to both Ctrl and Rescue animals. While bins corresponding to L4 are not significantly different, bins corresponding to L5b of the activated column and upper L2/3 of the neighboring column are higher in *Fmr1* KO animals and corrected in Rescue condition (p-values for Kruskal wallis test followed by [Tukey post hoc tests] for: Activated column, L4: Bin 25 p = 0.716, Bin 26 p= 0.444, Bin 27 p = 0.217, Bin 28 p = 0.108; L5b: Bin 42 p = 0.007 [Ctrl-KO p = 0.017, KO-Rescue p = 0.063, Ctrl-Rescue p = 0.999], Bin 43 p = 0.003 [Ctrl-KO p=0.012, KO-Rescue p = 0.033, Ctrl-Rescue p = 0.999], Bin 44 p = 0.016 [Ctrl-KO p = 0.062, KO-Rescue p = 0.039, Ctrl-Rescue p = 0.999]. Neighboring column, L2/3: Bin 6 p = 0.027 [Ctrl-KO p = 0.03263, KO-Rescue p = 0.190, Ctrl-Rescue p = 0.997], Bin 10 p = 0.073; L6: Bin 50 p = 0.049 [Ctrl-KO p=0.999, KO-Rescue p = 0.187, Ctrl-Rescue p =0.100], Bin 51 p = 0.037 [Ctrl-KO p = 0.999, KO-Rescue p = 0.146, Ctrl-Rescue p = 0.090]. **J**) Example of PV staining in S1: Scale bar: 40 μm. PV cell density, which is decreased in *Fmr1* KO mice, is restored in the Rescue condition. PV density Ctrl: 260.5 ± 6.7 N = 6, KO 225.9 ± 10.4 N = 6, Rescue 238 ± 5.9 N = 7; One-way ANOVA p = 0.022. Tukey post hoc tests: Ctrl/KO p = 0.019, Ctrl/Rescue p = 0.126, KO/Rescue p = 0.516. **K**) Example of cFos and PV colocalization. Scale bar: 40 μm. The ratio (%) of cFos^+^ PV neurons is increased in Fmr1 KO and restored in Early Rescue condition. Ctrl: 49.24 ± 5.90 N = 5; KO: 74.84 ± 2.97 N = 5; Rescue 46.96 ± 6.60 N = 6. One-way ANOVA p = 0.0071. Tukey post hoc tests: Ctrl/KO p = 0.026; Ctrl/Rescue p = 0.958; KO/Rescue p = 0.0085. **L**) Timeline of the novel texture discrimination assay and representation of Day 2 of the assay. 180-grit sandpapers are presented on two facing walls of the box during the Learning phase of the assay. 80-grit sandpaper (red) is presented as a novel textured object on one side, randomly during the Testing phase. Control and Rescue males can discriminate against the novel texture, while *Fmr1 KO* animals do not (Proportion of time spent exploring familiar *vs* novel texture - Ctrl: Familiar: 33 ± 5.3, Novel: 67 ± 5.3, p = 0.0085, N=13; *Fmr1 KO*: Familiar: 45.8 ± 4.2, Novel: 54.1 ± 4.2, p = 0.162, N=14; Rescue, Familiar: 33.8 ± 5.2, Novel: 66.2 ± 5.2, p = 0.024, N=10; One-sided Wilcoxon test for Novel > Familiar). N= animal, nSL = slice, n= cell replicates. Data are presented as mean ± SEM.

We next examined the previously described loss in firing precision of *Fmr1* KO upper-layer neurons^16,22^. In contrast to *Fmr1* KO neurons, the L4 neurons in the *Rescue* condition displayed a discrete firing pattern similar to the control neurons (Figure 3E). To assess the temporal gating of the L4 neuron firing, we examined the correlation between spikes within each condition (Figure 3F). Control L4 neurons formed discrete clusters of high correlation, while *Fmr1* KO animals exhibited a more continuous arrangement resulting in a higher mean correlation than control (Figure 3G). Notably, the clustering and the mean correlation of L4 neurons in the Rescue condition were more similar to those in control condition than Fmr1 KO animals. Bootstrap analysis further revealed that the higher mean correlation is induced only by *Fmr1* KO neurons, as they shift the mean correlation distributions of control and *Rescue* neurons closer to the *Fmr1* KO mean. In contrast, shuffling the control and *Rescue* datasets did not replicate the *Fmr1* KO-driven shift in correlation (Figure 3H).

We investigated whether the early SST neuron manipulation improves long-term neuron circuit integration within S1. We measured the immediate early gene *cFos* expression upon whisker activity from enriched environment exploration (Figure 3I, Figure S3E-F and methods)^35^. In this assay, animals with all whiskers trimmed except for the C1/C2, were allowed to explore an enriched environment for one hour. As expected, cFos intensity was the strongest in the sensory input recipient layers, L4 and L5b, of the cortical column corresponding to activated whiskers, in contrast to the neighboring columns corresponding to trimmed whiskers (Figure 3I; Figure S3G-H). Consistent with the FXS-associated hyperexcitability, cFos intensity was higher in *Fmr1* KO mice, although this increase was only significant in the bins corresponding to L5b, aligning with the results from L5 pyramidal neuron responses to TC inputs (Figure 3I, Figure S3H). Moreover, FXS-associated spatial diffusion into L2/3 of neighboring columns was also observed, consistent with previous studies (Figure 3I, Figure S3I)^36–38^. Strikingly, both features were corrected in the *Rescue* animals in addition to a signal decrease in L6 of the neighboring column.

Circuit dysfunctions in *Fmr1* KO animals have partially been attributed to PV neuron impairments^20,21,24,34^. We therefore next examined whether dysfunctions in PV neuron maturation occur downstream of persistent TC connectivity to SST neurons in *Fmr1* KO mice. Consistent with the previous studies, we observed a significant decrease in PV neuron density, specifically in L4 of *Fmr1* KO animals, yet this was not corrected by the *Rescue* strategy (Figure 3J; Figure S4A). While we found no anatomical defects in the number of PV synapses (Synaptotagmin2^+^/Gephyrin^+^/NeuN^+^) onto L4 neurons, which were previously described as hypoactive, presynaptic boutons were increased in *Fmr1* KO mice, suggesting a lack of terminal pruning. Notably, we observed a mild restoration in Rescue condition (Synaptotagmin2^+^/Gephyrin^+^/NeuN^+^; Figure S4B-C). In addition, the ratio of L4 PV neurons expressing cFos upon enriched environment exploration was increased in *Fmr1* KO mice and reduced to control levels in the *Rescue* condition (Figure 3K). This suggests that PV neuron integration into somatosensory circuits, but not their survival, is regulated by SST neuron maturation during postnatal stages.

The observation that SST neuron early dynamics control long-term circuit maturation in S1 suggests that they are involved in the establishment of adult tactile mouse behaviors. Altered tactile perception is known in children with FXS^18^ and previous studies revealed that mouse models of autism display behavioral sensory processing deficits^39,40^. In particular, *Fmr1* KO males lack whisker-based tactile discrimination behaviors^41,42^. Consistent with these studies, only control males significantly discriminated against distinct sand paper textures, while *Fmr1* KO animals did not exhibit this ability, using a similar whisker-based novel texture discrimination assay^43,44^. Remarkably, *Rescue* mice successfully discriminated between the textures in the task (Figure 3L and Figure S5).

Altogether, our results show that transient TC inputs to SST neurons during development are involved in downstream cortical maturation of sensory processing, specifically L4 excitatory and PV neuron maturation, as well as tactile mouse functions.

## Discussion

In this study, we unveil that transient dynamics between sensory thalamus and SST inhibitory neurons control downstream maturation of sensory functions. Moreover, we demonstrate that this connectivity is persistent, rather than dynamic, in a mouse model of FXS. Remarkably, restoring the transient dynamics during a specific postnatal critical period is sufficient to correct FXS-associated dysfunctions in somatosensory circuits and in sensory processing. Together, these findings provide the first direct evidence that perturbations in the transients of TC connectivity to SST neurons during development leads to FXS-linked defects.

Previous studies on FXS have focused largely on PV neurons rather than SST neurons. PV neurons constitute the feedforward motif of inhibition, which is triggered by sensory TC inputs in adulthood^1–3^ and defects in their function have been shown to contribute to not only neurodevelopmental disorders, but also psychiatric diseases^45^. Indeed, chemogenetic manipulations of PV neurons have been shown to restore both schizophrenia^46^- and FXS^20^-associated impairments. However, only late postnatal chemogenetic manipulation of PV neurons in *Fmr1* KO animals corrects defects in excitatory circuit^20^, while targeting all inhibitory circuits is successful at earlier postnatal stages^25,33^. Our results indicate that SST neurons are involved in early postnatal cortical circuit maturation and at earlier stages than PV neurons in neurodevelopmental disorders. Nonetheless, our early manipulation of SST neurons restored PV neuron circuit adaptation to sensory processing without affecting PV neuron density. This suggests that SST neurons contribute to specific, but not all aspects of PV neuron development. In *Fmr1* KO mice, PV neurons are expected to be hypoactive in response to evoked sensory stimuli^22^, suggesting that the increase in cFos expression in PV neurons and in the number of their boutons that we observed reflects their adaptive changes upon sensory processing impairments. In addition, we did not identify anatomical changes in PV neuron synapses between control and *Fmr1* KO animals, suggesting that SST neurons control intrinsic or other adaptive circuit properties of PV neurons rather than anatomical circuit rewiring. Thus, our findings provide insights into the mechanisms underlying the impaired maturation of PV neurons reported in FXS. Yet, further investigation of PV neuron circuit integration upon TC transient activation of SST neurons is warranted to dissect out PV neuron development downstream of SST neuron firing during development.

Abnormal sensory processing and hypersensitivity are critical symptoms of autism spectrum disorders, including FXS, and have also been reported in Fmr1 KO mice using sensory-dependent behavioral assays. Our findings using the novel texture discrimination task show that the persistent TC inputs to SST neurons play a key role in the processing of tactile information in adulthood. We have previously shown that SST neurons receive aberrant inputs from ventrobasal neurons thalamic population (VB), which is involved in passive whisking, as opposed to active whisking^12^. In the typical adult mouse brain, passive whisking is primarily integrated by PV neurons, while SST neurons shift from passive to active whisking during the second postnatal week^47,48^. In adulthood, PV neuron output mainly provides feedforward inhibition to L4 neurons, while SST neuron output constitutes L1-projecting feedforward inhibition. Therefore, our findings suggest that defects we observed in texture discrimination of FXS mice could result from improper feedforward inhibition from SST neurons receiving the aberrant VB information in addition to PV neurons in L4.

Beyond sensory TC inputs integration in L4, whisker-based sensory processing involves bottom-up and top-down circuit integration that occurs across distinct cortical layers, specifically in sensory-dependent behaviors. During the first postnatal weeks of development, SST neurons in L5 make transient translaminar contacts with L4-excitatory neurons^5,11,49^. They are thought to act as hub neurons in the cortex for the correlated-to-decorrelated shift in cortical activity^7,9,10^, as well as the shift from single- to multi-whisker sensory input integration in L2-3^48^. Therefore, these events potentially act as direct regulators of circuit corrections we observed downstream of transient connectivity restoration. The effect of the early *Rescue* strategy on the L5 pyramidal cell (inhibition and cFos expression) and L2/3 column organization, suggests the implication of SST neuron outputs in these layers. Interestingly, a recent study reported that SST neurons act as a time-gate and are necessary for callosal axon retraction from L4 neurons^50^. Interhemispheric connectivity is known to be reduced and the corpus callosum to be smaller in FXS patients and *Fmr1* KO animals^51–53^. Our findings reveal that SST neuron manipulation within a single hemisphere is sufficient for *Fmr1* KO animals to regain their texture discrimination abilities, suggesting that circuit restoration in the *Rescue* animals induced recovery of callosal projections. Further investigations into long-range cortical circuits would be highly valuable for elucidating the downstream consequences of transient restoration.

Lastly, a significant research objective in the exploration of neurodevelopmental disorders is to achieve substantial and lasting improvements in affected individuals through early therapeutic interventions^54^. Our results reveal that restoring the transient dynamics in TC input connectivity to SST neurons in S1 alleviates FXS-associated sensory impairments, suggesting that targeting transient connectivity onto SST neurons during postnatal development might be more effective in addressing autism-linked impairments. In line with this hypothesis, a recent study profiling single-cell transcriptional alterations in postmortem brains from autistic individuals revealed cortical SST neurons as one of the most transcriptionally altered populations in autism. Given that sensory hypersensitivity is an essential factor in autism-associated behaviors^17^, SST neurons could be a promising therapeutic target for early intervention in neurodevelopmental disorders.

## Acknowledgements

We thank Dr. Jessica Tollkuhn for sharing a camera and behavior boxes, Alister Orozco for producing supporting preliminary analyses, Dr. Dinu Albeanu for valuable advice on correlation analysis and Drs. Jessica Tollkuhn and Lucas Cheadle for critical reading of the manuscript.

This work was supported by a FRAXA fellowship to D.D, the National Institutes of Health (NIH) under Award Number R01MH136990 to GP. and under Award Numbers R01NS116897 and R01MH119818 to LVA. Additional funding support for this work is provided by the Simons Foundation (SFARI - Award: SFI-AN-AR-Pilot-00005618) and the Pershing Square Innovation Fund (PSIF) to G.P.

## Contributions

G.P. and D.D. conceived the project, developed the methodology, designed the figures and wrote the manuscript with inputs from S.L., S.G., L.H.L. and L.V.A.

G.P., D.D. and S.L analyzed and interpreted the results. D.D. performed electrophysiological experiments and corresponding analysis. S.L. and D.D. performed stereotactic surgeries. S.L. performed enriched environment experiments and generated all histology data. G.P. D.D. and S.L. performed synaptic/cellular quantification. D.V. participated in the quantification of cell density and colocalization. S.G., V.H.L, G.P.,D.D. and L.V.A designed the behavioral assay. S.G. and V.H.L performed behavioral assay recordings and video analyses. D.D. performed behavior statistical analysis.

## Declaration of interests

The authors declare no competing interests

## Methods

### Animals

All experiments were approved by and in accordance with Cold Spring Harbor Laboratories IACUC protocol number 22-4. Animals were housed under standard, temperature-controlled laboratory conditions and on a 12:12 light/dark cycle. Mice received water and food *ad libitum*. C57Bl/6 WT mice were used for breeding with transgenic mice. Transgenic mice: SST^Cre^ (stock number: 013044), *Fmr1* KO (stock number:003025) are available at Jackson Laboratories. Mice were injected at P0-P1 or at P20 for the Late Rescue strategy and experiments were conducted between ages P5-P6 for the early stage, P10-P16 for intermediate stage and between P30-P52 for the late time window. Both female and male animals were used in all experiments.

### Immunostaining

Mice were deeply anesthetized with sodium pentobarbital by intraperitoneal injection and transcardially perfused with PBS 1x followed by paraformaldehyde (PFA) dilutes at 4% in PBS. Brains were postfixed for 4 hrs in 4% PFA at 4°C. 50 µm vibratome (Leica) sections were incubated 1-2 hrs at room temperature in blocking solution containing 3% Normal Donkey serum and 0.3% Triton X-100 in PBS and incubated 48 hrs at 4°C with primary antibodies: Chicken anti-GFP (1:2000; Aves Labs #1020), Rabbit anti-RFP (1:500; Rockland #600-401-379), Mouse IgG1 anti-parvalbumin (1:500; Millipore #MAB1572), Rabbit anti-Somatostatin (1:1000; BMA Biomedicals #T-4103); Rabbit anti-FMRP (1:1000; Abcam #AB17722); Guinea Pig anti-cFos (1:1000; Synaptic Systems #226308), Guinea Pig anti-NeuN (1:500; Millipore #ABN90PMI), Mouse IgG2 anti-GAD65 (1:1000; Millipore #MAB351), Mouse IgG1 anti-Gephyrin (1:500; Synaptic Systems #147011), Mouse IgG2 Znp11 anti-Synaptotagmin 2 (1:500; ZFIN #ZDB-ATB-081002-25). Sections were rinsed three times 15 min with 0.1% Triton X-100 in PBS and incubated for 90-120 min at room temperature at room temperature with Alexa Fluor 488-, 555-, 647-conjugated donkey secondary antibodies (1:500; Invitrogen or Jackson ImmunoResearch) or DyLight 405 (1:500; Jackson Immunoresearch). Sections were mounted in Fluoromount-G (SouthernBiotech #0100-01) before imaging.

### Stereotactic injections

For postnatal timepoints, stereotaxic injections were performed using a neonate adapter (Harvard Apparatus; cat #75-1851). Mouse pups were anesthetized by hypothermia for 5 min, placed on a cooled and temperature monitored neonate adapter, cleaned with aseptic techniques and lidocaine 2% was applied to the injection site. Animals were then stereotaxically injected with AAV (volume 80-120nl) at P0-P1 with a nanoliter-injector (Nanoject III). For consistent targeting, cortical injections were performed over 5 min before pulling out the injector and at a rate of 1nL/sec. The primary somatosensory cortex was targeted with the following coordinates: AP: +1.40, ML: -1.70, DV: -0.20 from Lambda.

For the late rescue strategy, P20 mice received slow-release Buprenorphine subcutaneously before starting the surgery and were anesthetized in an isoflurane induction chamber at 5%, and then transferred in the stereotactic frame with 1-2% isoflurane. The primary somatosensory cortex was targeted using the following coordinates AP: +3.4 ML: -2.4, DV: -0.75 from Lambda.

AAV used in the study were as followed: AAV2-hSyn-DIO-hM3D(Gq)-mCherry (Titer: 6e-12 vg/mL) was a gift from Bryan Roth (Addgene plasmid #44361; http://n2t.net/addgene:44361; RRID:Addgene_44361)^55^ and AAV9-CAG-FLEX-EGFP-WPRE was a gift from Hongkui Zeng (Addgene plasmid # 51502; http://n2t.net/addgene:51502; RRID:Addgene_51502)^56^.

### Chemogenetics for Early and Late Rescue strategies

Clozapine N-oxide (CNO) dichloride (HelloBio, HB6149) was dissolved in 0.9% saline at 1mg/mL (10x solution) and stored at 4°C for the duration of the chronic activation (up to 10 days). Neonates were weighted each day for subcutaneous injection of CNO at 1 mg/kg once a day. For the early rescue strategy, CNO was injected from P6 to experiment time (from P10 to 16) to examine the intermediate stage and from P6 to P16 to examine the late stage. For the late rescue strategy, animals received CNO from P26 to P36.

Toxicity and specificity of AAV-infection was verified by immunostaining of Ctrl, KO and Rescue animals with somatostatin. 1-2 slices per animal, with the injection site labeled, were imaged at 10x using a Zeiss epifluorescence microscope. Ratio of reporter^+^ AAV infected cells expressing somatostatin was quantified using Fiji (ImageJ) software in selected S1 region, across all layers. On the same slices, somatostatin^+^ cells were quantified in a defined region of interest (ROI), to measure density of SST neurons across conditions.

### Enriched environment and cFos expression assay

Adult (P48-52) mice were anesthetized with 5% isoflurane in an induction chamber. Once fully unconscious, they were placed under a microscope under light anesthesia: 0.5-1% isoflurane. The entire right whisker pad was trimmed using microscissors, while all but C1 and C2 whiskers were trimmed on the left side. After whisker trimming, mice were allowed to fully recover from anesthesia before being placed in a cage enriched with multiple novel objects, in a quiet and dark room. Mice were perfused after 1 hour spent exploring the cage.

Mouse slices were stained for cFos. For each animal, the section with the most visible cFos^+^ L4 barrel corresponding to C1/C2 whisker activation was selected for cFos expression intensity analysis. Intensity measured throughout the activated column (through L4-cFos barrel) and throughout the neighboring column, from the top, Layer 1 to SVZ level, using “Plot Profile” in Fiji software. Average pixel intensity was measured with distance (in pixels). To normalized expression levels, background was defined as the minimum value measured within L1 (pixel 2-60), which lacked cell bodies/nuclei and therefore cFos expression, and it was subtracted from each value. L1-6 distance in pixels was organized into ∼50-51 bins, centered from the highest L4-value in the activated column, and from the highest L2 (pixels 2-11) value in the neighboring barrel. Data distribution was considered not normal based on the Kolmogorov-Smirnov test and therefore non-parametric, Kruskal-Wallis test was performed on Bins with possible *Fmr1* KO-associated effects (highlighted in gray) and followed by Dunn’’s post hoc multiple comparison test between each condition.

Colocalization of cFos expression within PV neurons was manually identified using Fiji Cell counter, from 40x confocal images of L4 and L5 within the activated column and of L4 within the neighboring column. The percentage of PV neurons expressing cFos out of all visible PV labeled cells was measured within each image

### Synaptic contact analysis

PV neuron output was identified through immunofluorescence of Synaptotagmin 2 (presynaptic boutons), Gephyrin (postsynaptic density) onto L4 neurons labeled with NeuN. On a separate set of staining, SST neuron output was identified from AAV-infected SST neurons expressing a cytosolic reporter, GAD65 (presynaptic inhibitory marker) and Gephryin (postsynaptic density) onto L4 neurons labeled with NeuN.

Number of synaptic contacts or puncta was quantified similarly to previously published synaptic analysis^12^. L4 NeuN^+^ soma were located within the injection site, and ∼10 cells from 3 different brain slices were imaged for each animal. For more homogeneous cell replicates, one plan was imaged for each cell. The centers of soma were identified when the NeuN signal had the highest diameter. A LSM 780 Zeiss confocal was used to image the cells with a 63x oil objective, a 6x digital zoom, at 2048x2048 pixels, 16-bit resolution. Two different experimenters performed the imaging and analysis in order to maintain the experimenter #2 blind to the analysis. Experimenter #1 next re-assigned each blinded quantification to the animal and condition for statistical analysis.

Kolmogorov-Smirnov tests identified PV neuron presynaptic boutons as normal and all other measurements as non-normal. Therefore one-way ANOVA, followed by Tukey’s post hoc tests and Kruskal-Wallis, followed by Dunn’s post hoc tests were applied, respectively.

### PV neuron density

PV neuron density across layers was measured on the same experiments used for enriched environment assays. PV labeling was imaged from the cFos-labeled single “activated” slices per animal at 10x using a Zeiss epifluorescence microscope. Fiji (ImageJ) software was used to define a Region of Interest (ROI) throughout the S1 “activated” column, in which all PV labeled neurons were manually counted. Layer-specific PV density was quantified from 40x images using a Zeiss LSM800 confocal microscope in L4 and L5 of the S1 activated column. The same images were used for quantification of colocalization of cFos^+^ PV neurons.

### Electrophysiology

#### Thalamocortical slices

Mice from P5 to P40 were anesthetized with isoflurane and decapitated. Brains were dissect in 4°C artificial cerebro-spinal fluid (aCSF) containing (in mM): NaCl 125, KCl 2.5, NaH2PO4.H2O 1.25, NaHCO3 25, D-Glucose 20, CaCl2 2, MgCl2 1, pH adjusted to 7.4 with NaOH. Thalamocortical (TC) slices from P5 to P40 were obtained by using a guide to cut the brains at specific angles as previously described^2,27,28^. P10-16 brains were placed on an angled platform to create a 10° angle with the horizontal plan. Then, the brains were cut at ⅓ of their antero-posterior length from the anterior part with a 50° angle from the midline. For P30-40 brains, TC slices were made as above with a 55-60° angle with the midline. P5-6 brains were pre-embedded in 3% low melt agarose made in cutting aCSF containing (in mM): NaCl 87, KCl 2.5, NaH2P04.H20 1.25, NaHCO3 26, Sucrose 75, D-Glucose 10, CaCl2 1, MgCl2 2, ph adjusted to 7.4 with NaOH. Then, the embedded brains were flipped to the side, centered on a xy frame and positioned in such a way that the most anterior-ventral and posterior points touched respectively the x and the y axis. Then, a 20-25° angle from the x axis was made in the agarose block, a few millimeters below the anterior-ventral part. Then, embedded brains could be tilted at 20-25° and received a second cut at 55° from the midline as for P10-40. Brains were sliced with a vibratome (Leica, VT-1000S) in 4°C carbonated (95% O2, 5% CO2) cutting aCSF and placed in carbonated aCSF at room temperature for 2-3 minutes to slowly re-equilibrate the ions before being kept at 37°C in carbonated aCSF.

#### Recordings

TC slices from P5 to P40 were placed in the recording chamber of a SliceScope pro 1000 (Scientifica) constantly perfused with a carbonated aCSF at 4 ml per min. Perfused aCSF was warmed up by a heater (TC-324C, Warner instrument) to reach 35-37 in the recording chamber. Electrodes were pulled from filamented thick-wall borosilicate glass (B150-86-7.5, Sutter instrument) with a vertical puller (P-30, Sutter instrument). Brain slices illuminated by an infra-red light source were visualized with a camera (SciCam Pro, Scientifica) through a 20X immersion objective (XLUMPLFLN20X, Olympus). Brain slices were screened and only those with conserved TC fibers were used. TC fibers were electrically stimulated with a stimulator generator (model 2100, A-M systems) whose positive electrode was placed in the recording chamber and the negative one was inserted in a 1-2 MΩ borosilicate glass pipette filled with an HEPES-based aCSF containing (in mM): NaCl 125, KCl 2.5, CaCl2 2, NaH2PO4.H2O 1.25, HEPES 10, MgCl2 1, pH adjusted to 7.4 with NaOH. Stimulation pipettes were placed in the middle of the TC fibers beam exiting the thalamus. Slices were exposed to 470 or 550 nm light from a fluorescence LED source (PE-4000, CoolLED) to visualized SST neurons infected by AAVs.

SST neurons in S1 L5 were then recorded in whole cell configuration of the patch clamp technique, and held at -70 mV in voltage clamp mode with a 5-6 MΩ borosilicate glass pipette filled with an internal solution containing (in mM): Cs-Methanesulfonate 124, NaCl 10, EGTA 1, HEPES 10, ATP-Mg 4, GTP-Na 0,4, pH adjusted to 7.4 with CsOH, osmolarity adjusted to ∼290 mOSM with Cs-Methanesulfonate.

At P10-16, TC EPSCs were recorded from L5 Pyramidal neurons in the vicinity of connected SST neurons (Figure S1 C-D) or not (All Pyramidal neurons in Figure S2 F-J) as described above. GABAergic currents evoked poly-synaptically by TC electrical stimulation were recorded at 0 mV. Pyramidal neurons were identified by their large soma.

Excitability properties of L5 SST neurons in S1 were recorded in whole cell configuration and current clamp mode with a 5-6 MΩ borosilicate pipette filled with an internal solution containing (mM): K-Methanesulfonate 125, NaCl 10, EGTA 1, HEPES 10, ATP-Mg 4, GTP-Na 0.4, 1% biocytin, pH adjusted to 7.4 with NaOH, osmolarity adjusted to ∼290 mOSM with K-Methanesulfonate. Forty increment steps of 10 pA were injected from an initial step = (holding current -100 pA).

Action potentials (AP) from S1 L5 SST neurons were recorded in loose cell attach configuration with a 5-6 MΩ borosilicate glass pipette filled with the same HEPES-based aCSF used for electrical stimulation. All SST neurons recorded in this way were localized near TC-connected SST cells and the TC fibers were stimulated the exact same way as to evoked EPSCs.

Putative L4 excitatory neurons were identified by their soma size and recorded in loose cell attached configuration as described above. Layer 4 boundaries were estimated by the SST cell density and the distance to Layer 1. AP were elicited by single electrical stimulation of the subventricular zone (SVZ). Bath applications of aCSF containing NBQX (2uM) + APV (50uM) or GABAzine (5uM) were used to control the synaptic origin of the AP evoked by SVZ stimulation (all drugs from Hello Bio).

For whole cell recordings, the pipette capacitance was canceled after the formation of the giga-seal and the cell one was canceled 1-2 min after breaking in. Cells with an access > 30 MΩ were not concidered for recording. The series resistance was compensated to 40-60%. Evoked currents and AP were sampled at 20 KHz and filtered at 3 and 8.5 KHz respectively with an amplifier (EPC10, HEKA). Each recording contains between 100 to 200 trails.

#### Biocytin staining of L4 neuron

L4 neurons identified as described above were recorded in whole cell configuration with a borosilicate pipette filled with an internal solution containing (mM): K-Methanesulfonate 125, NaCl 10, EGTA 1, HEPES 10, ATP-Mg 4, GTP-Na 0.4, 1% biocytin, pH adjusted to 7.4 with NaOH, osmolarity adjusted to ∼290 mOSM with K-Methanesulfonate. After 10 minutes from breaking the membrane, the pipette is slowly removed to reach the outside-out configuration. Slices were fixed 1-2 hours in PFA 4%, washed twice 15 min in PBS at RT and left O/N in PBS at 4°C. Slices were then incubated 24 hours at 4°C in PBS containing 0.6% Triton-X (Sigma), 0.3% cold fish gelatin (Sigma) and 1:1000 Streptavidin antibody conjugated with an Alexa fluor 555 (Thermofisher #S21381). Before mounting, slices were washed out three times 15 minutes in PBS. Filled L4 neurons were imaged with a fluorescence microscope and a 20x dry objective.

#### TC connectivity analysis

TC fibers stimulation trials that did not evoke EPSCs or IPSCs were not included in the analysis due to the impossibility to assign these events to stimulation or transmission failure. Postsynaptic current peak amplitude and charge were measured from averaged traces. Cells not responding to the electrical stimulation of the TC fibers were not included in this study.

#### L5 SST neurons firing analysis

SST neurons not responding to the electrical stimulation of the TC fibers were included when neighboring ones were found to be connected with TC fibers. The firing probability was calculated as the proportion of trials with AP in a 50 ms time window from the stimulation onset. *L4 neurons firing analysis:* Cells not responding to the electrical stimulation of the SVZ fibers were not included in this study. Firing probability was calculated as the proportion of trials without AP. L4 neurons exhibiting more than 60 % of action potential failure were not added to the analysis of firing patterns and correlation matrices. The firing probability was calculated as the proportion of AP in 1 ms bins. Pearson correlations were computed between every pair of neurons per condition and sorted to optimize their clustering. *Bootstrap analysis:* Ctrl and Rescue data are pooled and 72 neurons (the *Fmr1* KO group size) are randomly drawn with replacement from this pooled distribution. Next, a correlation matrix is generated as above and the mean correlation is measured. This process is repeated 10K times, generating the ‘Ctrl + Rescue’ distribution. To test whether neurons from the *Fmr1* KO group yield more variability, the same process is repeated from “Ctrl + *Fmr1* KO” and “Rescue + *Fmr1* KO” pooled dataset. P-values were calculated as the probability of each mean correlation distribution to be equal or above the observed mean correlation from the *Fmr1* KO L4 neurons.

#### SST neurons excitability analysis

Input resistance (IR) was estimated as the slope of the linear regression between the net injected current and the mean potential from the sweeps before firing occurs. Firing current was estimated as the first current step evoking more than one AP. AP threshold (th) was estimated as the potential corresponding to the maximal second derivative of the first AP from the first sweep with more than one AP. Resting membrane potential (RMP) was estimated as the potential corresponding to the holding current on the linear regression between the mean potential and the net injected current from the sweeps before firing occurs.

### Whisker-based novel Texture Discrimination task

P40-P53 animals were subjected to a whisker-based novel texture discrimination task as previously reported ^43,44^. The testing setup consisted of a 33 cm x 33 cm white acrylic chamber which was placed inside a box with an infrared camera mounted to the top. The box served to block extraneous olfactory, visual, or audio stimuli. A day prior to testing, mice were habituated to the testing arena for 10 minutes. The next day, each mouse was subjected to three phases: habituation, learning, and test. During the habituation phase, the mouse was allowed to explore the empty testing chamber for 5 minutes and then removed to a separate cage for 5 minutes. During the learning phase, fine 180 grit sandpaper strips (9 cm x 7 cm) were affixed to the center bottom of opposite walls and the mouse was permitted to explore the chamber for 5 minutes before being removed to the separate cage for 1 minute. During this time, the sandpaper strips were replaced with a fresh 180 sandpaper strip (familiar object), and a coarse 80 grit sandpaper strip (novel object). Subsequently in the test phase, mice were returned to the chamber for 3 minutes. Video from the infrared camera was analyzed in EthoVision XT 11.5 (Noldus). The mouse was defined as investigating a texture when its nose point was located at most 2 cm away from the texture. The amount of time the mouse spent investigating the textures was measured for the learning and testing phases. The proportion of time spent exploring textures is defined as the time spent investigating a texture normalized by the total time spent investigating either texture. The position of the novel texture (right or left side) during the testing phase was randomized and did not impact mouse performance (Spearman correlation between time spent exploring side of the novel texture during learning (%) and time spent exploring the Novel texture during testing (%); Ctrl: 0.44, p=0.128; KO: 0.11, p=0.68; Rescue: -0.44, p=0.197). Animals who spent less than 1 second interacting directly with one of the two textures were excluded from the analysis. Using these criteria, 1 female Rescue, 5 females and 2 males *Fmr1* KO, 3 females and 2 males Ctrl were excluded. The chamber was cleaned with 70% ethanol between mice to eliminate olfactory cues.

## Notes

### Competing Interest Statement

The authors have declared no competing interest.

