## Supplementary figures and table for "Restoring transient connectivity during development improves dysfunctions in fragile X mice"

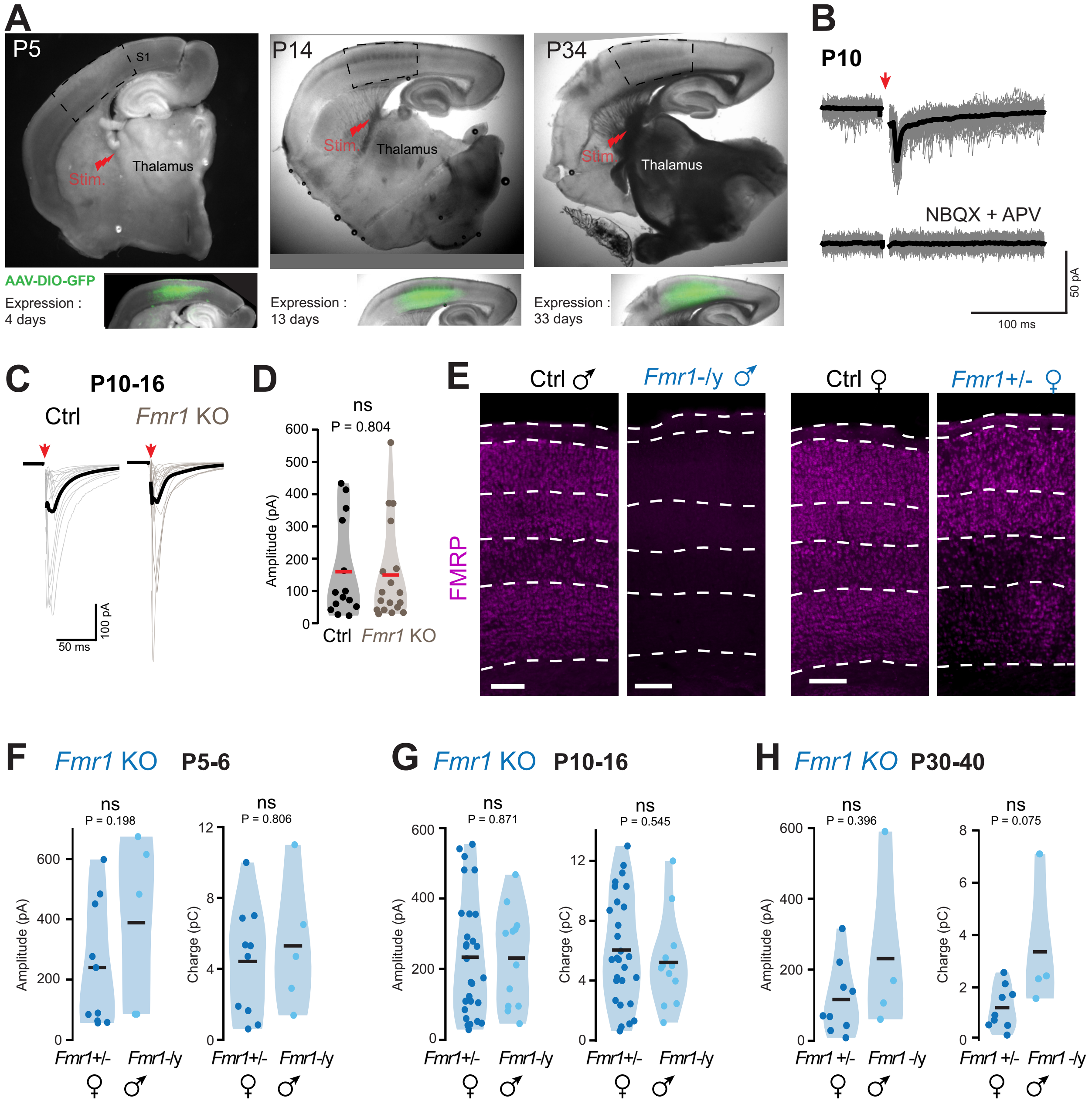


**Figure S1. TC connectivity in *Fmr1* KO mice, Related to Figure 1**

**Figure S1: TC connectivity in *Fmr1* KO mice, Related to Figure 1.**
**A**) Example of TC slice for each time window, P5 for the early window, P14 for the intermediate (same example as Figure 1) and P34 for the late time window. Bottom: AAV-DIO-GFP expression in S1. Stim = electrical stimulation of TC fibers. **B**) Single EPSCs (gray) and average (black) from a single SST neuron, evoked by TC fiber stimulation. The response is abolished by glutamatergic transmission blockers, NBQX (2uM) and APV (50uM). **C**) Traces of every averaged EPSCs evoked in L5 pyramidal cells by TC electrical stimulation at P10-16. **D**) Amplitude of the averaged EPSCs evoked in L5 neurons show no significant difference between control and *Fmr1* KO mice (Amplitude: Ctrl 159.66 ± 39 pA, n = 14; KO 149.37 ± 35 pA, n = 18; Charge (data not plotted) Ctrl: 5.89 ± 1.17 pC; KO : 3.94 ± 0.69 pC; Bilateral Mann Whitney U test p = 0.106). **E**) Example of FMRP staining showing the absence of FMRP expression in *Fmr1* KO hemizygous males (*Fmr1*-/y) and the mosaic, reduced expression in heterozygous females (*Fmr1*-/+). **F-H**) The distribution of average EPSC amplitude and charge is not specific to males or females within the *Fmr1* KO animals, during the early P5-6 (F), intermediate P10-16 (G) and late P30-40 window (H) [p-values from Bilateral Mann Whitney U test]. (P5-6 Amplitude, F: 239.62 ± 61.6 pA, n=10; M: 388.4 ± 114.2 pA, n=5; p = 0.198; Charge, F: 4.4 ± 0.93 pC; M: 5.3 ± 1.5 pC; p = 0.806; P10-16 Amplitude, F: 233.2 ± 31.28 pA, n=28; M: 230.8 ± 38.19, n=12; p = 0.871; Charge, F: 6.04 ± 0.7 pC; M: 5.21 ± 0.8 pC; p = 0.545; P30-40 Amplitude, F: 115.23 ± 32 pA, n=9; M: 230.83 ± 105.5 pA, n=4; p = 0.396; Charge, F: 1.22 ± 0.25 pC; M: 3.36 ± 1.09 pC; p = 0.075). N= animal, nSL = slice, n= cell replicates. Data are presented as mean ± SEM.

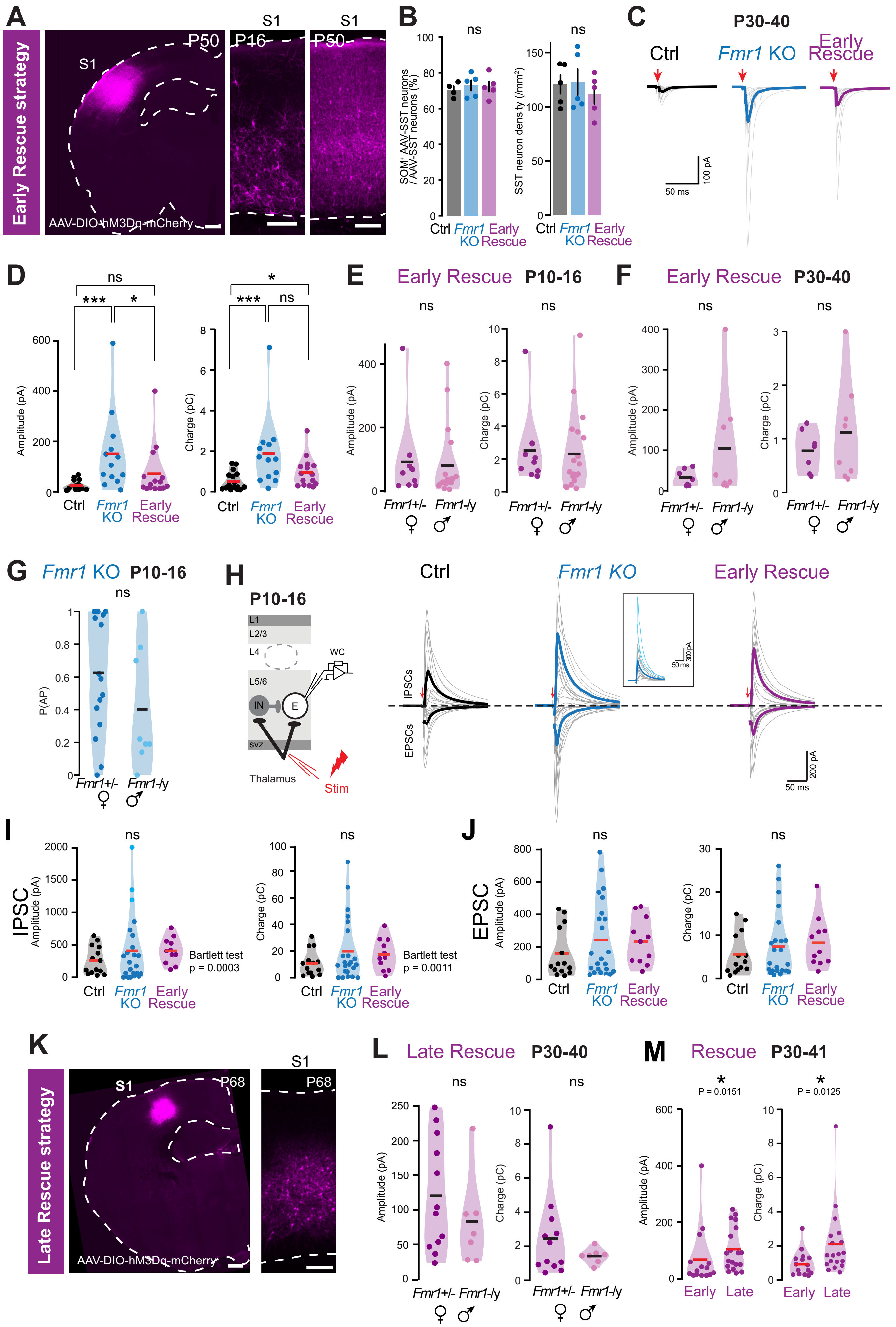


**Figure S2. Rescue strategy: TC inputs onto SST neurons, Related to Figure 2**

**Figure S2: TC connectivity in *Fmr1* KO mice, Related to Figure 1.**
**A**) Example injection site of AAV-DIO-hM3Dq-mCherry for Early Rescue at P50 at low magnification. Scale bar: 500 µm. Left inset: example of injection in S1 at P16, Right inset: injection site in S1 at P50 from the lower magnification image. Scale bar: 200 µm. **B**) Absence of toxicity and specificity of AAV-infection across conditions: Percentage of AAV-infected cells identified as somatostatin positive ($SOM^+$) neurons. (Ctrl 70.29 ± 2.03 % N = 4, *Fmr1* KO 72.74 ± 2.84 % N = 5, Rescue 72.15 ± 2.81 % N = 5). $Somatostatin^+$ SST neuron density (cell number/$mm^2$) (Ctrl: 120.7 ± 8.530 N = 5, KO: 122.7 ± 12.02 N = 5, Rescue: 111.4 ± 8.474 N = 5; One-way ANOVA: p = 0.69). **C**) Traces of average EPSCs evoked in Ctrl, *Fmr1* KO and Early Rescue SST neurons by TC fiber stimulation during the late time window, P30-40. During the late time window, TC inputs are maintained stronger onto *Fmr1* KO SST neurons compared to control neurons. **D**) Amplitude and charge of average EPSCs at P30-40. Ctrl and *Fmr1* KO EPSC data are from Figure 1. (Amplitude: Ctrl 24 ± 4.4 pA, n = 17, nSL = 4, N = 4; KO 150.8 ± 42 pA, n = 13, nSL = 4, N = 1M/3F; Rescue 71 ± 26 pA, n=15, nSL = 4, N = 2M/2F; Kruskal-Wallis test p =0.0013 , Dunn's post hoc tests: Ctrl/KO p = 0.00028, KO/Rescue p = 0.042, Ctrl/Rescue p = 0.10; Charge: Ctrl 0.51 ± 0.1 pC; KO 1.88 ± 0.46 pC; Rescue 0.95 ± 0.18 pC; Kruskal-Wallis test p=0.0013, Dunn's post hoc tests: Ctrl/KO p = 0.00032, KO/Rescue p = 0.125, Ctrl/Rescue p = 0.0352). **E-F**) The distribution of averaged EPSC amplitude and charge is similar across Early Rescue males and females, during the intermediate P10-16 and the late P30-40 time window (P10-16 Amplitude, F: 92.32 ± 38.8 pA, n=10; M: 79.59 ± 24.6 pA, n=19; p = 0.476; Charge, F: 2.53 ± 0.67 pC; M: 2.31 ± 0.53 pC; p = 0.435; P30-40 Amplitude, F: 32.22 ± 7.11 pA, n=7; M: 104.94 ± 45.1 pA, n=8; p = 0.602; Charge, F: 0.77 ± 0.13 pC; M: 1.11 ± 0.31 pC; p = 0.684; Bilateral Mann Whitney U test). **G**) The distribution of *Fmr1* KO SST neuron firing probability is not significantly different across sexes at P10-16 (F: 0.62 ± 0.08, n=16; M: 0.4 ± 0.12, n=8; Bilateral Mann Whitney U test p = 0.166). **H**) Schematic illustration of L5-excitatory neuron whole-cell recordings and traces of average EPSCs and IPSCs in Ctrl, *Fmr1* KO and Early Rescue animals. Inset: blue traces show highest IPSCs not represented in the main panel. **I**) Average IPSC evoked in L5-excitatory neurons by TC fiber stimulation. Amplitude Ctrl: 259 ± 55.7 pA, n=14; KO: 410.6 ± 105.7 pA, n=23; Rescue: 410 ± 57.8 pA, n=11; Kruskal-Wallis test p = 0.236; Bartlett test p = 0.00033; Charge, Ctrl: 10.7 ± 2.45 pC; KO: 19.7 ± 4.8 pC; Rescue: 17.3 ± 3.38 pC; Kruskal-Wallis test p = 0.346, Bartlett test p = 0.0011). **J**) Average EPSC evoked in L5-excitatory neurons by TC fiber stimulation (Amplitude Ctrl: 159.8 ± 38.87 pA; KO: 243.25 ± 47.9 pA; Rescue: 233.7 ± 41.5 pA; Kruskal-Wallis test p = 0.346, Bartlett test p = 0.103; Charge, Ctrl: 5.56 ± 1.22 pC; KO: 7.38 ± 1.53 pC; Rescue: 8.28 ± 1.66 pC; Kruskal-Wallis test p = 0.346, Bartlett test p = 0.204). **K**) Example injection site of AAV-DIO-hM3Dq-mCherry for Late Rescue at P68 at low magnification. Scale bar: 500 µm. Inset: injection site in S1. Scale bar: 200 µm. **L**) The distribution of average EPSC amplitude and charge is not significantly different across Late Rescue males and females (Amplitude, F: 82.74± 22.94 pA, n=7; M: 120.27 ± 22.37 pA, n=12; Bilateral Mann Whitney U test p = 0.374; Charge, F: 1.43 ± 0.15 pC; M: 2.45 ± 0.66 pC; Bilateral Mann Whitney U test p =0.735). **M**) Average EPSC amplitude and charge of SST neurons within the Early Rescue is higher than with the Late rescue strategy at P30-40. Bilateral Mann Whitney test (Amplitude E/L p = 0.015; Charge E/L p = 0.012). N= animal, nSL = slice, n= cell replicates. Data are presented as mean ± SEM.

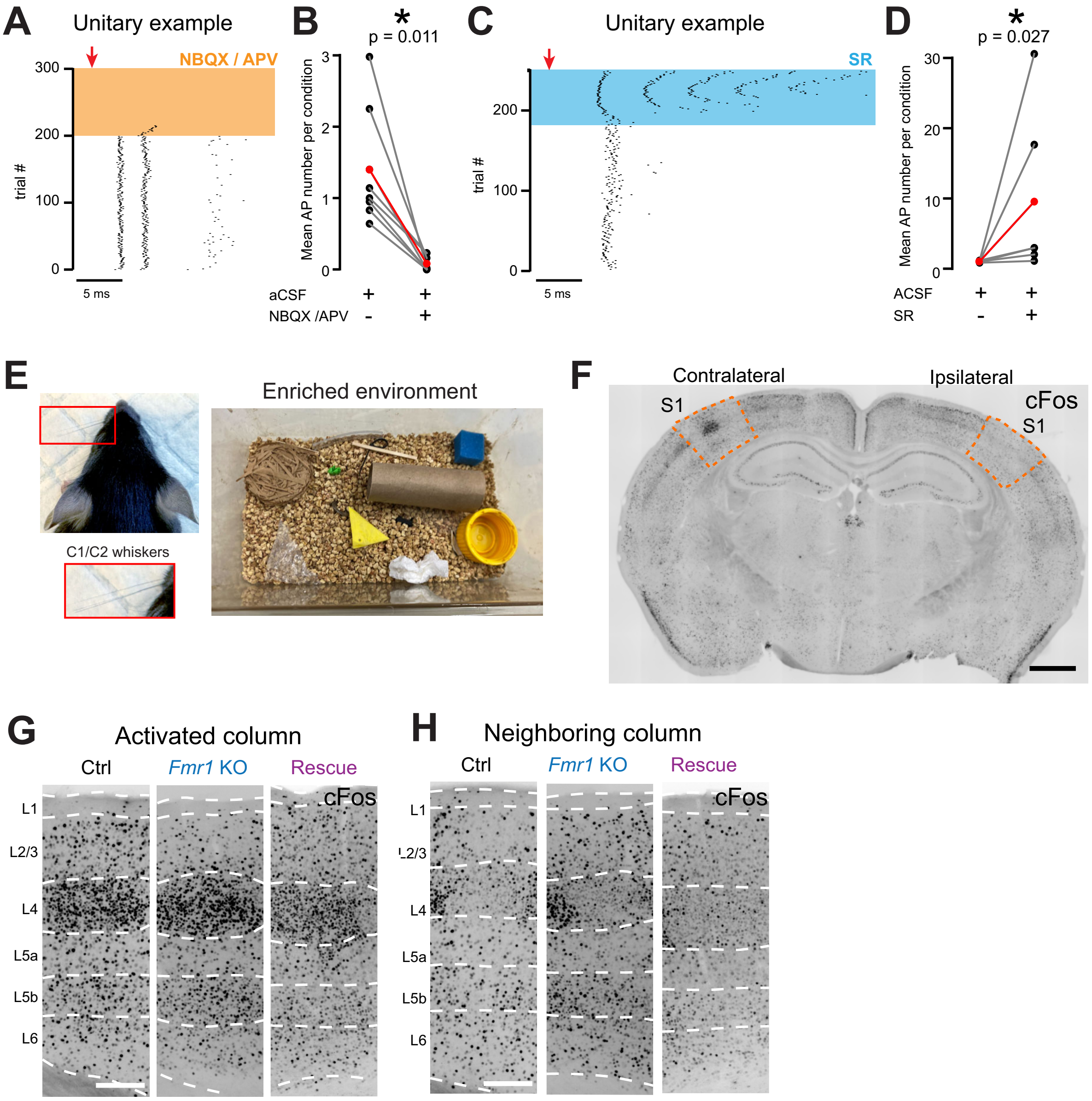


**Figure S3. Somatosensory circuit function in *Fmr1* KO and Rescue animals, Related to Figure 3**

**Figure S3: Somatosensory circuit function in Fmr1 KO and Rescue animals, Related to Figure 3.**

**A**) Unitary example of a L4 neuron firing evoked by electrical stimulation of the subventricular zone (SVZ, red arrow) recorded before and after application of glutamatergic transmission blockers: NBQX (2 µM) and APV (50 µM) in orange. **B**) Mean of action potential (AP) numbers per condition (aCSF) and in presence of glutamatergic blockers (aCSF: 1.39 ± 0.26, aCSF+NBQX+APV: 0.081 ± 0.035, n=8, nSL=8, N= 4 *Fmr1 KO* & 4 Ctrl, Bilateral Wilcoxon paired test p = 0.011) Average in red. **C**) Same as A) but in presence of GABAergic blocker: SR (10 µM) in blue. **D**) Same as B) in presence of SR (aCSF: 1.02 ± 0.04, aCSF+SR: 9.54 ± 4.48, n=6, nSL=6, N= 1 *Fmr1 KO* & 5 Ctrl, Bilateral Wilcoxon paired test p = 0.027). **E**) Example of the trimming of all whiskers, but the C1 and C2 whiskers on one side, before enriched environment exploration. Image of the enriched environment. **F**) cFos expression from both S1 control (ipsilateral) and C1/C2 activated (contralateral) sides. Scale bar: 1mm. **G-H**) cFos staining in the activated (G) and the neighboring (H) columns of a Ctrl, *Fmr1* KO and Rescue animal. Scale bar: 200 µm.N= animal, nSL = slice, n= cell replicates. Data are presented as mean ± SEM.

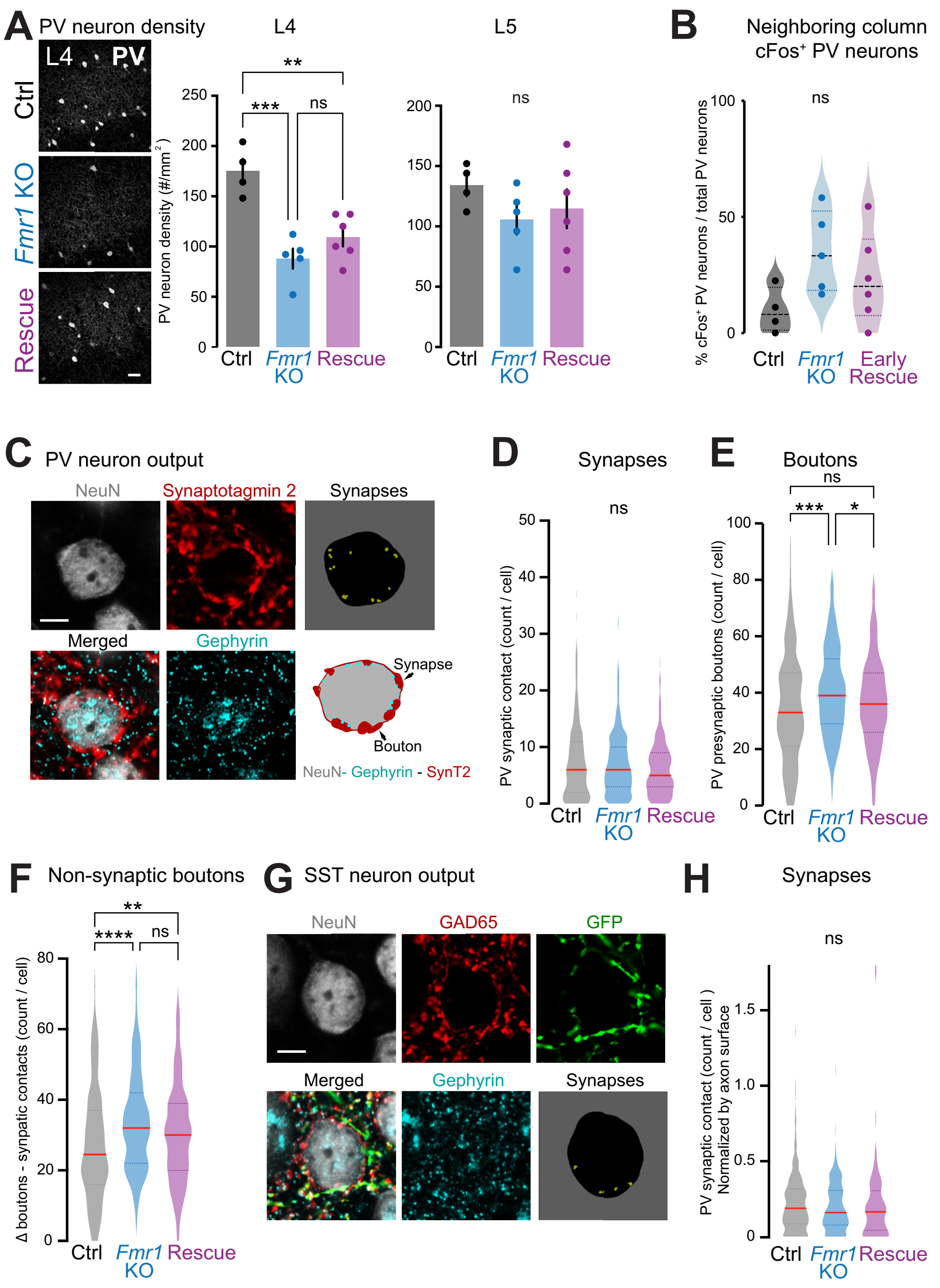


**Figure S4. PV neuron development in *Fmr1* KO and Rescue animals, Related to Figure 3**

**Figure S4: PV neuron development in *Fmr1* KO and Rescue animals, Related to Figure 3.**

**A**) PV density in Ctrl, *Fmr1* KO and Rescue L4. Same image as Figure 3K. Scale bar: 40 µm. PV density in L4 (Ctrl 49.24 ± 5.90 N = 4, KO 74.84 ± 2.97 N = 5, Rescue 49.96 ± 6.60 N = 6, One-way ANOVA p = 0.007, Tukey post hoc tests: Ctrl/KO p = 0.026, KO/Rescue p = 0.008, Ctrl/Rescue p = 0.958). PV density in L5 (Ctrl 124.3 ± 13.07 N = 4, KO 117.6 ± 5.48 N = 5, Rescue 102 ± 10.21 N = 6, One-way ANOVA p = 0.287). **B**) As expected from the lack of direct whisker activation, the ratio of cFos$^+$ PV neurons in the L4 of the neighboring column is not significantly different across conditions. Ctrl: 9.67 ± 4.87 N = 4; KO: 35.00 ± 7.89 N = 5; Rescue 23.41 ± 7.95 N = 6. One-way ANOVA p = 0.124. **C**) Images of L4 neurons (NeuN+) labeled with PV neuron output markers: Synaptotagmin 2 (SyT2) = presynaptic boutons, Gephyrin = postsynaptic density. Mask of automated synpase detection. Scale bar: 5 µm. **D**) Number of PV neuron synaptic contacts in L4 (SyT2$^+$+Gephyrin$^+$) per cell (NeuN$^+$) is similar across Ctrl, *Fmr1* KO and Rescue animals. (Ctrl: 7.82 ± 0.36 n = 330 N = 4F/5M; KO: 6.90 ± 0.38 n = 205 N = 2F/3M; Rescue: 6.19 ± 0.25 n = 253 N = 4F/3M; Kruskal-Wallis test p = 0.479). **E**) Number of PV neuron boutons (SyT$^+$) in L4 (NeuN$^+$) from the same analysis exhibit more presynaptic puncta in *Fmr1* KO mice compared to Ctrl mice. PV neuron boutons are not significantly different from both Ctrl and KO animals. (Ctrl: 34.48 ± 1.04; KO: 40.53 ± 1.07; Rescue: 36.64 ± 0.98; One-Way ANOVA test p = 0.0004, Dunn's post hoc tests: Ctrl/KO p = 0.0002, KO/Rescue p = 0.041, Ctrl/Rescue p = 0.28). **F**) Number of non-synaptic boutons (SyT2$^+$ - (SyT$^+$+Gephyrin$^+$)) is increased in both *Fmr1* KO and Rescue L4 compared to Ctrl L4. (Ctrl: 27.20 ± 0.87; KO: 33.62 ± 0.95; Rescue: 30.45 ± 0.86; Kruskal-Wallis test p < 0.0001, Dunn's post hoc tests: Ctrl/KO p < 0.0001, KO/Rescue p = 0.079, Ctrl/Rescue p = 0.0091). **G**) Images of L4 neurons (NeuN$^+$) labeled with SST neuron output markers: AAV-DIO-GFP$^+$ + GAD65$^+$= presynaptic boutons, Gephyrin = postsynaptic density. Mask of automated synpase detection. Scale bar: 5 µm. **H**) Number of SST neuron outputs (GFP$^+$+GAD65$^+$+Gephyrin$^+$) normalized by GFP$^+$ axonal surface, to L4 neurons (NeuN$^+$) is not significantly different between Ctrl, *Fmr1* KO and Rescue mice. (Ctrl: 0.226 ± 0.013 n = 236 N = 3F/4M; KO: 0.197 ± 0.011 n = 213 N = 2F/4M; Rescue: 0.238 ± 0.042 n = 117 N = 1F/3M (Kruskal-Wallis test p = 0.215). N = animal, nSL = slice, n= cell replicates. Data are presented as mean ± SEM.

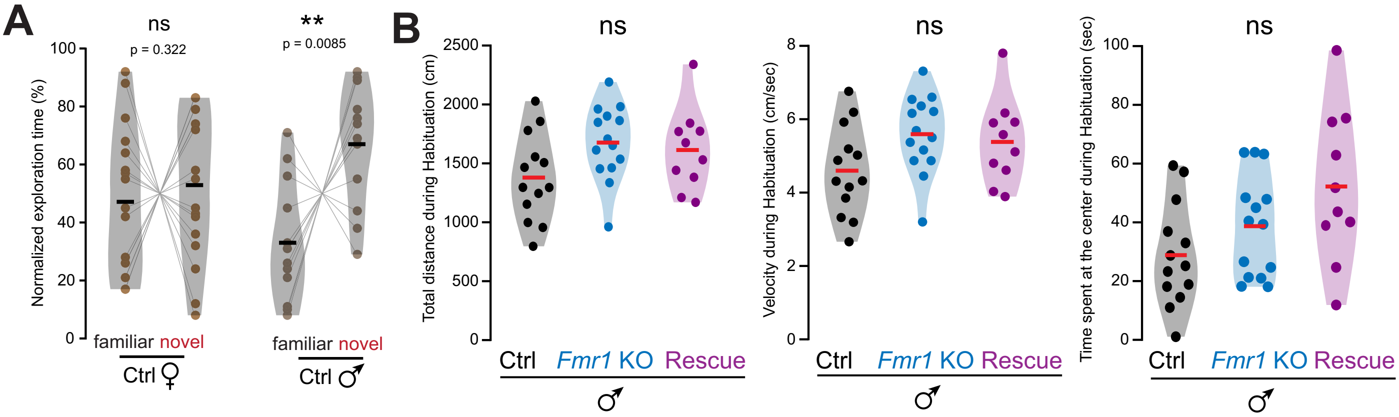


**Figure S5. Whisker-based texture discrimination assay, Related to Figure 3**

**Figure S5: Whisker-based texture discrimination assay, Related to Figure 3.**
**A**) Ctrl females discrimination is not displayed in the novel texture discrimination assay. (Proportion of time spent exploring familiar *vs* novel texture; Ctrl, Familiar: 47.1 ± 5.8, Novel: 52.8 ± 5.8, p = 0.322, N = 17; One-sided Wilcoxon test for Novel > Familiar). **B-D**) No motor and anxiety defects are identified across Ctrl, *Fmr1* KO and Rescue animals. B) Total distance traveled by males during Habituation (Ctrl: 1379.1 ± 97.61 cm; KO: 1676 ± 81.46 cm; Rescue: 1613.1 ± 104.1; Kruskal-Wallis test p = 0.096). C) Velocity of males during Habituation (Ctrl: 4.59 ± 0.32 cm/sec; KO: 5.59 ± 0.27 cm/sec; Rescue: 5.38 ± 0.38 cm/sec; Kruskal-Wallis test p = 0.097). D) Time spent at the center of the open field during Habituation, Ctrl: 28.83 ± 4.7 sec; KO: 38.69 ± 4.45 sec; Rescue: 52.2 ± 7.78 sec; Kruskal-Wallis test p = 0.062). N= animal, nSL = slice, n= cell replicates. Data are presented as mean ± SEM.

| | Mean ± SEM | | | Kruskal-Wallis | Dunn's post hoc tests (p-values) | | |
|---|---|---|---|---|---|---|---|
| | Ctrl | Fmr1KO | Rescue | (p-values) | Ctrl vs KO | Ctrl vs Rescue | KO vs Rescue |
| Input Resistance (Mohm) | 480.3 ± 51 | 242.7 ± 17 | 255.75 ± 32 | 0.0010 | 0.0011 | 0.0014 | 0.99 |
| Firing current (pA) | 39.1 ± 6 | 87.7 ± 13 | 94.16 ± 16.5 | 0.010 | 0.0073 | 0.0097 | 0.96 |
| AP threshold (mV) | -54.5 ± 1,57 | -52.4 ± 1,8 | -53.9 ± 2.26 | 0.83 | - | - | - |
| RMP (mV) | -70.5 ± 1.06 | -73 ± 0.74 | -70.4 ± 0.37 | 0.018 | 0.0090 | 0.71 | 0.029 |
| n | 12 | 13 | 12 | | | | |
| nSL | 2 | 4 | 3 | | | | |
| N | 1M/1F | 2M/2F | 1M/2F | | | | |

**Table S1. Intrinsic properties of SST neurons in Ctrl, *Fmr1* KO and Rescue mice, Related to Figure 2.**
